# Long-term epidemiology and evolution of swine influenza viruses in Vietnam

**DOI:** 10.1101/2023.02.08.527780

**Authors:** Jonathan Cheung, Anh Ngoc Bui, Sonia Younas, Kimberly M. Edwards, Huy Quang Nguyen, Ngoc Thi Pham, Vuong Nghia Bui, Malik Peiris, Vijaykrishna Dhanasekaran

**Author notes:** These senior authors contributed equally to this article (Malik Peiris, Vijaykrishna Dhanasekaran).

## Abstract

Swine influenza virus (SwIV) surveillance in Hanoi, Vietnam from 2013–2019 found gene pool enrichment from imported swine from Asia and North America. Long-term maintenance, persistence and reassortment of SwIV lineages was observed. Co-circulation of H1-δ1a viruses with other SwIV genotypes raises concern due to its zoonotic potential.

## Introduction

Influenza A viruses (IAVs) circulate in multiple host species and pose a persistent One Health threat because of their ability to spill over into new host populations (1). As avian and mammalian viruses can infect pigs and undergo reassortment, pigs are considered to play an important role as “mixing vessels” in the generation of viruses with pandemic potential (2). The 2009 H1N1 pandemic was caused by a reassortant H1N1 virus containing gene segments from classical swine virus, Eurasian avian-like swine virus (EA), and human seasonal virus lineages (3). Three different IAV subtypes (H1N1, H3N2, and H1N2) circulate in swine worldwide (4), with regional variation in antigenic characteristics. Contemporary swine influenza virus (SwIV) lineages include H1N1, H1N2, and H3N2 triple reassortant viruses (TR) that originated through reassortment of classical swine, avian, and human influenza viruses in the 1990s (1, 5), European avian-like (EA) H1N1 SwIV introduced into pigs during the 1970s, and several human H1N1 and H3N2 derived viruses, including H1N1pdm09, pre-2009 H1N1 and H3N2-variant viruses (4). The increasing genetic diversity of IAVs in swine presents a pandemic concern (4, 6). Human infections of emerging SwIV are most frequently reported from the U.S.A. because all non-human influenza viruses are nationally notifiable in the U.S.A., with outbreaks occurring in agricultural fairs (7, 8), however sporadic human cases due to SwIVs have recently been reported in Asia (9, 10), Europe (11), and Australia (12).

In 2013, SwIV surveillance was established at a collective slaughterhouse in Hanoi, Vietnam that sourced pigs from 23 local provinces (13) (**Table 1**). During 2013–2014, co-circulation of multiple SwIVs was detected, including viruses originating from H1N1pdm09, H1N2 with pre-2009 seasonal influenza-derived H1 hemagglutinin (HA), H3N2 derived from human seasonal influenza, and ‘triple-reassortant’ (TR) H1N2 and H3N2 viruses (13). Here, we extend data on virological surveillance to 2019 and provide data on serological surveillance from 2013 to 2019 in this slaughterhouse to characterise SwIV evolution. Genome sequencing showed continuous enrichment of H1 and H3 subtype diversity through repeat introduction of human virus variants and SwIVs endemic in other countries. In particular, the North American H1-*δ*1a with a TR backbone that facilitates increased human adaptation, may pose a zoonotic threat.

**Table 1.** Summary of swine influenza virus isolates by province.

| Province of origin | No. samples | No. (%) SwIV positive |
| --- | --- | --- |
| Ha Noi | 2178 | 52 (2.39)* |
| Ha Nam | 855 | 10 (1.17) |
| Hai Phong | 61 | 7 (11.48) |
| Bac Ninh | 51 | 4 (7.84) |
| Thai Nguyen | 199 | 3 (1.51) |
| Ninh Binh | 22 | 1 (4.55) |
| Nghe An | 51 | 1 (1.96) |
| Hung Yen | 96 | 1 (1.04) |
| Vinh Phuc | 215 | 1 (0.47) |
| Hoa Binh | 465 | 2 (0.43) |
| Phu Tho | 301 | 1 (0.33) |
| Bac Giang | 200 | 0 (0) |
| Yen Bai | 150 | 0 (0) |
| Thanh Hoa | 45 | 0 (0) |
| Thua Thien Hue | 31 | 0 (0) |
| Nam Dinh | 29 | 0 (0) |
| Dien Bien | 2 | 0 (0) |
| Thai Binh | 135 | 0 (0) |
| Tuyen Quang | 27 | 0 (0) |
| Dong Nai, SV | 195 | 11 (5.64) |
| Binh Duong, SV | 16 | 0 (0) |
| Binh Dinh, SV | 10 | 0 (0) |
| Kieu Ninh, SV | 7 | 0 (0) |
| Location not specified | 509 | 20 (3.93) |
| **Total** | **5,850** | **114\* (1.95)** |

\* Includes one mixed infection of H1N2 and H3N2: A/swine/Hanoi/9-179/2017
† SV, southern Vietnam

## Results

### SwIV incidence and subtype diversity

One hundred and fifty paired serum and nasal swab specimens were collected monthly during 2016–2019 through convenience sampling at the Van Phuc slaughterhouse, the main collective slaughterhouse in Hanoi, Northern Vietnam. SwIVs and subtypes detected are shown in **Fig 1a** and **Fig 1b**. For ease of comparison, virus detection and subtypes from previous studies conducted in Vietnam from 2010 to 2016 have been included in **Fig 1**. No clear pattern of seasonality was observed (**Fig 1a**). The abattoir sourced swine from 23 provinces in Vietnam (**Fig 1c**). The majority of samples (87%) originated from provinces in northern Vietnam, while 4% of samples originated from southern provinces. Location data was missing for the remaining samples (9%). Most SwIV positives originated from Hanoi (46%), Ha Nam (9%), and Hai Phong (6%) in Northern Vietnam, and Dong Nai (10%) in Southern Vietnam (**Fig 1c, Table 1**). There were 114 influenza positive nasal swabs with one mixed-subtype infection among the 5,850 swabs tested, yielding 115 viruses for study, with an isolation rate of 1.95%.

**Figure 1.**
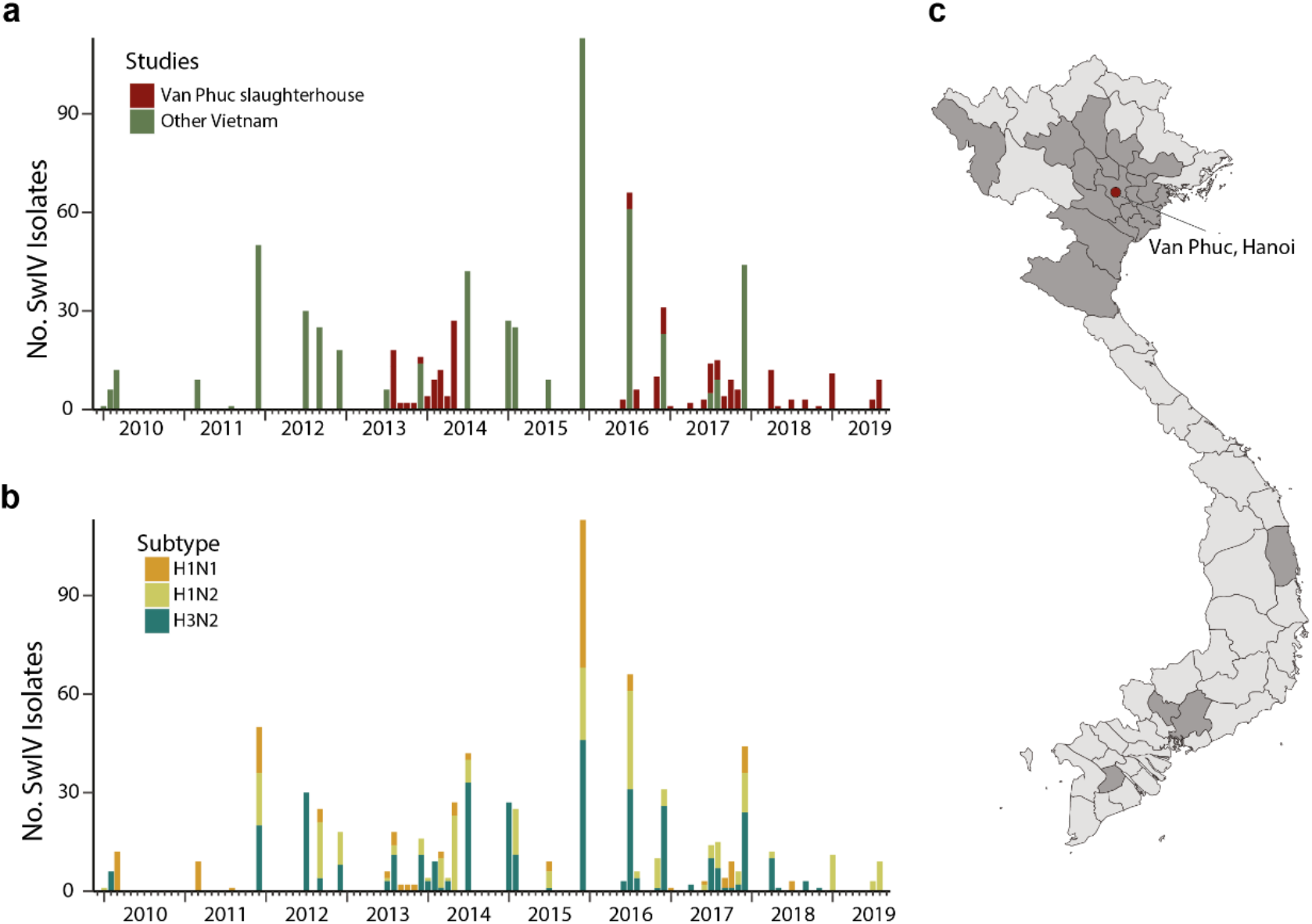
Swine influenza virus detection in Vietnam between 2010–2019. (a) Numbers of swine Influenza viruses isolated from Van Phuc slaughterhouse during 2013–2014 (13, 14) and 2016–2019 (denoted in red), alongside other surveillance studies in Vietnam shown in green (15, 16, 17). (b) Prevalence of swine influenza virus subtypes. (c) Provincial origins of pigs sampled at Van Phuc slaughterhouse, Hanoi, during 2016–2019.

Subtyping of the 115 SwIV isolates collected from June 2016 to August 2019 showed the circulation of three virus subtypes, H1N1 (n=16), H1N2 (n=46), and H3N2 (n=53), including one specimen with a mixed H1N2 and H3N2 infection. Our previous study in the same abattoir from 2013–2014 also yielded the same three virus subtypes, H1N1 (n=16), H1N2 (n=35) and H3N2 (n=26) (13). Notably, we observed a high genetic similarity of viruses collected during each sampling at the Van Phuc slaughterhouse, suggesting that viruses were not maintained in the abattoir, but were repeatedly introduced from sourced populations.

### Phylogenetic relationships of the HA genes

**Fig 2a** shows a maximum likelihood (ML) phylogenetic analysis of the HA genes of 145 SwIVs collected from 2016–2019 in Vietnam, together with virus sequences from previous studies from the same abattoir during 2013–2019, and 390 SwIVs collected in Vietnam by other studies. HA sequence data reveals the circulation of four distinct H1 lineages and two distinct H3 lineages over various time points (**Fig 2b**, Fig S1). The H1 subtype viruses include human-derived H1N1pdm09 (n=122), classical swine-derived H1-TR (n=84), and pre-2009 seasonal H1N1 (n=70) which are further classified into H1-δ-like (n=60) and H1-δ1a (n=9). Additionally, two viruses of the H1N2 subtype collected in December 2016 by Takemae et *al*. (2018) (17) belong to the Eurasian avian (EA)-like swine H1 HA lineage. The H3-HA phylogeny indicates SwIV in Vietnam belong to either the swine-origin North American TR lineage or the 2003/04 human H3N2-origin SwIV first detected in Vietnam in 2010 (**Fig 2b**, Fig S1).

**Figure 2.**
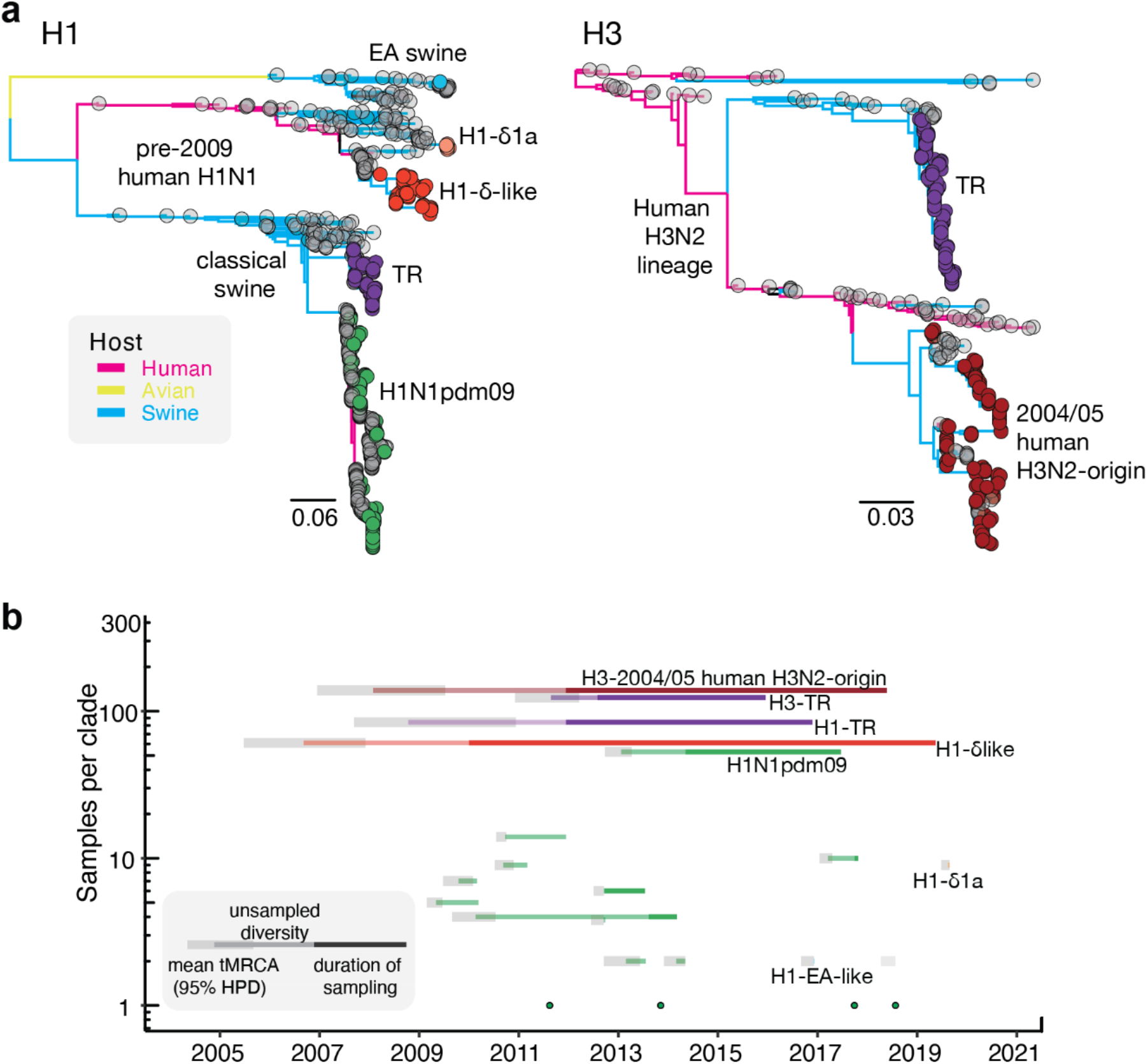
Genomic epidemiology of swine influenza viruses in Vietnam. (a) Maximum likelihood phylogeny of the H1 (left) and H3 (right) hemagglutinin (HA) genes of swine influenza viruses (SwIVs) in Vietnam, with tips colored by lineage origin and branches colored by host. (b) Persistence of independent HA lineages of SwIVs detected in Vietnam. H1N1pdm09 lineages are shown in green. TR, triple reassortant; EA, Eurasian avian.

HA phylogenies identified the circulation of 20 independent transmission lineages in swine in Vietnam since 2010, including four singletons (**Fig 2b**). These lineages were all identified elsewhere in the world and were likely introduced by imported swine or by humans through reverse zoonosis. Of the 20 transmission lineages, 16 independent transmission lineages of the H1N1pdm09 virus were detected at various time points. Most introductions were limited to 1–14 samples collected during one to three contiguous sampling months and were most closely related to spillover of contemporary human H1N1pdm09 viruses. However, two closely related swine H1N1pdm09 viruses that appear to have independently derived from humans around 2013 were detected in January and June 2017 (A/swine/Hanoi/7_619/2017 and A/swine/Hanoi/8_463/2017), indicating persistence of H1N1pdm09 for more than three years in swine. In addition to recurrent detections of H1N1pdm09 between 2014–2017 (n=53); a single lineage of H1 TR virus (n=84), most closely related to viruses collected from swine in Guangxi during 2010–2011, was maintained from 2010–2019; and an H1-δ-like lineage (n=60) derived independently from the pre-2009 human seasonal H1N1 viruses that circulated in humans around 2004–2006. In addition, nine virus sequences from August 2019 collected on the last sampling occasion belonged to H1-*δ*1a lineage and were most closely related to H1N2 SwIVs circulating in swine in the U.S.A. during 2015–2016. The H1-*δ*1a originated from human seasonal H1N1 viruses circulating in humans during 2002/03 (18, 19), and was previously only found in swine in the U.S.A. (20).

Two major H3-HA lineages originated from North America. One belongs to the TR lineage that circulated from 2012–2015 in Northern Vietnam (n=132) (13, 14, 16), and the other belongs to the 2004/05 human H3N2-origin SwIV (n=133) that circulated from 2010–2018 (**Fig 2b**). A zoonotic case reported in Vietnam during 2010 (A/Ho Chi Minh/459-6/2010 (H3N2)) and SwIV collected in southern China and Cambodia during 2011–2012 clustered with the 2004/05 human H3N2-origin SwIV. The 2004/05 human H3N2-origin viruses have been further classified into two divergent lineages, NV and SV, based on detection in north and south Vietnam, respectively. SV viruses have not been detected since 2015, and NV viruses were detected up to 2018. Taken together, our results indicate that the overall HA diversity of SwIV in Vietnam may have reduced in recent years, with H1 and H3 TR lineages not detected since 2016 and detections of new human H1N1pdm09 viruses declining in recent years. However, H1-*δ* viruses became predominant by 2018/19.

Temporal phylogenies of the H1 and H3 HA genes of Vietnam SwIV (**Fig 2b** and **Fig 3a**) identified long-branches of unsampled diversity leading up to the H1-*δ*-like virus and the 2004/05 human H3N2-origin SwIV, which was prior to initiation of swine surveillance in Vietnam, as well as the H1N1pdm09 virus lineage detected four years after it was first detected in 2013 (described above). Long-branches of unsampled diversity spanning four to five years were also identified within the Vietnamese SwIV clades. A sublineage of the 2004/05 human-origin swine virus in Vietnam which diverged before 2010 was only detected in this study between 2016–2019. Similarly, recent viruses belonging to two H1-*δ* sublineages appear to have been sampled after four years.

**Figure 3.**
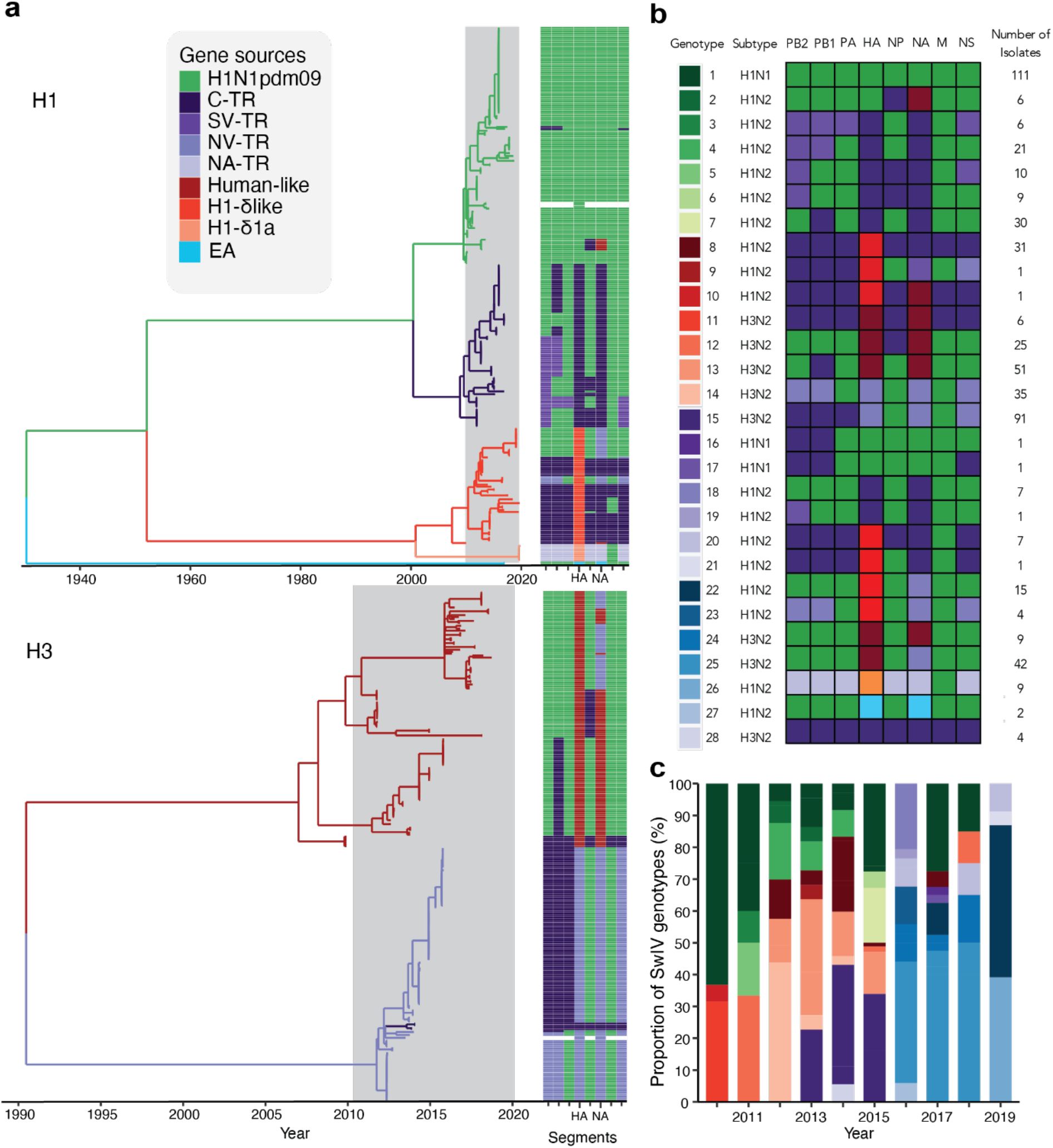
H1 and H3 swine influenza virus genotypes in Vietnam, 2010–2019. (a) Temporal phylogeny of the HA genes of SwIVs isolated in Vietnam. Branch colors denote virus lineage from which HA gene segments originate. Heatmap illustrates gene segment origins. C-TR, Chinese triple reassortant; SV-TR, Southern Vietnam triple reassortant; NV-TR, Northern Vietnam triple reassortant; NA-TR, North American triple reassortant; BD-like, Binh Duong-like human seasonal H3; H1-*δ*, pre-2009-human H1-*δ*; H1-*δ*1a, pre-2009-human H1-*δ*1*a*; EA, Eurasian avian. (b) Genotypes identified in Vietnam from 2010-2019. Complete SwIVs sequences and genotype numbers defined by Takemae et *al*. (16) and newly detected genotypes. Box colors denote the virus lineage from which the gene segment originates. Green, H1N1pdm09; dark purple, Chinese triple reassortant (C-TR); indigo, Southern Vietnam triple reassortant (SV-TR) cluster; lilac, Northern Vietnam triple reassortant (NV-TR); mauve, North American triple reassortant (NA-TR); maroon, human seasonal H3-like; red, H1-δ-like; orange, H1-δ1a; cyan, Eurasian-avian (EA)-like.

### SwIV genotypes in Vietnam 2010–2019

Phylogenetic analysis of each gene-segment of SwIVs from 2010–2019 showed the SwIV gene pool in Vietnam contains genes from H1N1pdm09 virus, pre-2009-human derived H1 viruses further subdivided into H1-*δ*-like and H1-*δ*1a, EA and TR viruses of H1 and H3 subtype (**Fig 3**). Based on the origins of individual gene segments, SwIV in Vietnam up to 2015 were previously designated into genotypes 1–15 (13, 16). We detected an additional 13 new genotypes in Vietnam between 2016–2019, which we designate as genotypes 16–28. Three of the original 15 genotypes (1, 8, and 12), as reported in (13, 16), continued to be detected during 2016–2019. Genotypes 27 and 28 were reported elsewhere (23, 20) respectively.

In each year we observed the co-circulation of multiple genotypes, with few genotypes detected across multiple years (**Fig 3c**). However, predominant genotypes associated with each of the major H1 and H3 HA lineages circulated during various periods. The majority of SwIV detected in Vietnam were genotype 1, with all eight gene segments originating from human H1N1pdm09 viruses (n=111) (**Fig 3**, S1). Genotype 1 was dominant in Vietnam from 2010–2015 and 2017–2018, while H1N1pdm09 viruses with PB2, PB1, NP and NS genes from TR viruses and N2-NA genes from human H3N2-derived SwIV (genotypes 2, 16, and 17) were detected sporadically (**Fig 3**). Similar to the HA gene, the NA and internal genes of swine H1N1pdm09 viruses were also not monophyletic but interspersed between human-origin virus sequences. All of the potential spillover events were closely related to human viruses, not limited to human viruses in Vietnam.

In contrast to the predominance of a single H1N1pdm09 virus genotype, the H1N2 TR viruses repeatedly gained internal genes from H1N1pdm09 viruses (genotypes 3–7, 18–19), with the most recent H1N2 TR viruses containing all internal genes of H1N1pdm09 viruses. Two H1N2 SwIV sequences (genotype 27) contain HA and NA genes of EA-like origin and all internal genes from H1N1pdm09. Similarly, the genotype 8 H1N2 viruses, with H1-*δ* HAs and all other segments from TR viruses, repeatedly gained H1N1pdm09 virus genes to form genotypes 20–23 and 26. The most recent H1N2 TR viruses collected in 2019 contain TR N2 and H1N1pdm09 internal segments (genotype 22). However, H1-*δ* genotype 8 viruses with no H1N1pdm09 genes continued to be detected until 2019. Furthermore, H1-*δ*1a viruses (genotype 26) isolated in August 2019 contained North American TR internal and NA genes with only the MP gene from H1N1pdm09. These viruses have not been seen anywhere other than the U.S.A. and now, Vietnam, suggesting that these SwIVs likely spread to Vietnam directly from the U.S.A.

The H3 subtype SwIV in Vietnam formed eight genotypes. The H3 TR viruses that circulated in north Vietnam between 2012–2015 were initially detected with all genes of TR lineage (genotype 28) or with PA, NP and MP genes derived from H1N1pdm09 viruses (genotype 14). However, H3 TR viruses detected during 2014– 2015 contained only NP and MP genes from H1N1pdm09 while maintaining the remaining TR genes (genotype 15). The 2004/05 human H3N2-derived viruses, initially identified in 2010 with all internal genes from TR lineages (genotype 11), gained H1N1pdm09 gene segments with or without maintaining TR PB1 (genotype 13) or NP genes (genotype 12). However, most recently collected viruses of this H3 lineage in 2017–2018 contained the TR N2 gene and all six internal genes of H1N1pdm09 (genotype 25).

Taken together, SwIV lineages of all subtypes collected in recent years in Vietnam have acquired the H1N1pdm09 internal genes, except two divergent H1-*δ* viruses that appear to have become predominant in Vietnam in recent years.

### Seroprevalence of SwIV in northern Vietnam

From 2013 to 2019, ten sera were randomly selected from each abattoir visit for serological analysis through hemagglutination inhibition (HAI) assays. Of swine sera tested, 41% (309/760) showed HAI antibody titers ≥40 against at least one of five representative virus antigens tested (**Table 2**). Seroprevalence of 2004/05 human H3N2-origin virus in 2016–2019 (15.3%) was broadly comparable to 2013–2014 (15.9%), but changes were seen in H1 lineages. Seroprevalence of H1N1pdm09 and H1-TR virus decreased, while H1-*δ*-like virus seroprevalence increased from 13% in 2013–2014 to 20% in 2016–2019. H1-*δ*1a antibodies were only detected in 2016– 2019, with the first seropositive samples collected in March 2016 (Table S1) and had limited antibody cross-reactivity to other H1 SwIV lineages (Table S2). Despite continuous detection of an H1-TR lineage between 2013–2017 (**Fig 2b**), overall seroprevalence to H1-TR reference strain (A/swine/Hanoi/7_305/2016(H1N2)) remained low at 2.4% (**Table 2**). H1-TR seropositive samples displayed elevated HAI titers for H1N1pdm09 (2–3 fold increase, Table S3). This suggests that H1-TR reactive antibodies are likely due to cross-reactivity with H1N1pdm09.

**Table 2.**
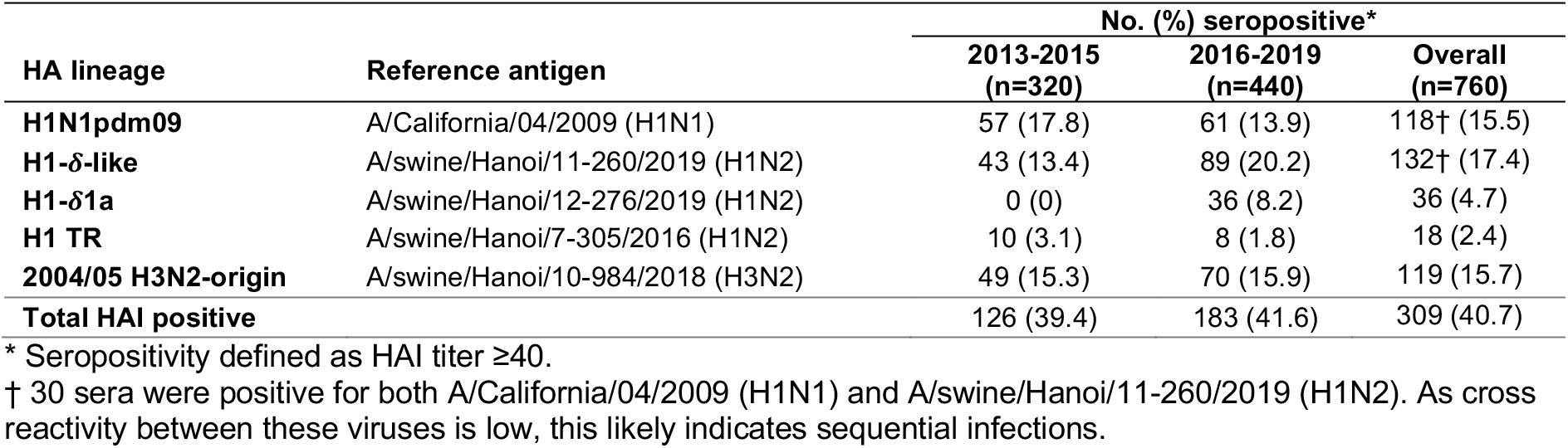
Hemagglutination inhibition (HAI) seroprevalence against different lineages of influenza A viruses circulating in Northern Vietnam

| HA lineage | Reference antigen | No. (%) seropositive* |  |  |
| --- | --- | --- | --- | --- |
|  |  | 2013-2015<br>(n=320) | 2016-2019<br>(n=440) | Overall<br>(n=760) |
| H1N1pdm09 | A/California/04/2009 (H1N1) | 57 (17.8) | 61 (13.9) | 118† (15.5) |
| H1-δ-like | A/swine/Hanoi/11-260/2019 (H1N2) | 43 (13.4) | 89 (20.2) | 132† (17.4) |
| H1-δ1a | A/swine/Hanoi/12-276/2019 (H1N2) | 0 (0) | 36 (8.2) | 36 (4.7) |
| H1 TR | A/swine/Hanoi/7-305/2016 (H1N2) | 10 (3.1) | 8 (1.8) | 18 (2.4) |
| 2004/05 H3N2-origin | A/swine/Hanoi/10-984/2018 (H3N2) | 49 (15.3) | 70 (15.9) | 119 (15.7) |
| **Total HAI positive** |  | 126 (39.4) | 183 (41.6) | 309 (40.7) |

\* Seropositivity defined as HAI titer $\geq 40$ .
† 30 sera were positive for both A/California/04/2009 (H1N1) and A/swine/Hanoi/11-260/2019 (H1N2). As cross reactivity between these viruses is low, this likely indicates sequential infections.

## Discussion

Our study, conducted from 2016–2019, provides insights into the evolution and epidemiology of SwIV in Vietnam, a major pork-producing country in Asia. Through continuous surveillance at a central slaughterhouse in Hanoi, which sourced pigs from across the country, we found that only H1N1, H1N2 and H3N2 were present and co-circulating. This is consistent with previous surveillance at the same slaughterhouse in 2013/14 (13, 14) and other studies in Vietnam (15, 16, 17), indicating that the SwIV subtypes in Vietnam are similar to those in other countries in Asia and other continents (21, 22, 23, 24).

Our analyses of SwIV sequence data from Vietnam since 2010 showed that the genetic diversity was due to the persistence of several swine-origin H1N2 and H3N2 viruses, reported first in other countries in Asia and North America and imported into Vietnam, likely via swine trade. We also identified repeat introductions of human H1N1pdm09 viruses through reverse zoonosis. We found extensive reassortment of the SwIV internal genes among the major SwIV lineages circulating in Vietnam, with H1N2 and H3N2 SwIV gaining internal genes from H1N1pdm09 viruses but only rare acquisition of other SwIV internal genes by H1N1pdm09 lineages. As a result, most recent Vietnamese swine viruses contain H1N1pdm09 internal genes. However, two divergent H1-*δ* viruses (originally derived from pre-2009 seasonal H1N1 viruses), detected in our most recent sample from 2019, maintained the TR lineage NA and internal genes (genotypes 20, 21 and 26, **Fig 3b**).

While repeat introduction of H1N1pdm09 into swine is consistent with previous swine surveillance data from Vietnam (13) and southern China (25, 26), the frequency of detections of new human H1N1pdm09 virus lineages in swine in Vietnam has reduced since 2015. Although onward transmission of H1N1pdm09 was not sustained for most introductions, one lineage, first detected in northern Vietnam during 2013/14 (13), continued to circulate in swine for more than four years and reassorted with other prevailing SwIV lineages. Similar evidence of H1N1pdm09 being maintained within swine populations has been reported from the U.S.A. (27) and Australia (28).

Pre-2009 seasonal H1N1 influenza-derived swine viruses (classified under H1-*δ*) are gaining predominance in swine in Vietnam. Notably, an H1N2 H1-*δ*1a virus lineage with limited cross-reaction to other H1 SwIV was detected in 2019, and serology showed evidence of circulation since March 2016, indicating a continual enrichment of SwIV from other regions. The increasing prevalence of H1-*δ*-like lineages increases the risk for zoonotic transmission as children and adolescents may have little or no immunity to pre-2009 seasonal H1 viruses. Previous studies demonstrated significant antigenic distance between H1N1pdm09-like viruses and H1-*δ* cluster (H1-*δ*-like and H1-*δ*1-a) (29, 30) and current seasonal influenza vaccines do not elicit protection against H1-*δ*-like viruses. To identify such zoonotic transmission, it is crucial to ascertain whether current diagnostics can detect and distinguish pre-2009 H1 viruses from currently circulating human seasonal H1 viruses.

Furthermore, H1-*δ*1a viruses likely entered Vietnam via imported swine, demonstrating the role of trade in the global dissemination of SwIVs (31). East and Southeast Asian countries including China, Thailand, and Vietnam, which produce more than 50% of pork production globally, are hotspots for emerging infectious diseases from swine (32, 33). Vietnam ranks second in Asia for pork production, producing 42.2 million heads in 2019 (34). Pork consumption in Vietnam risen rapidly, from 12.8 kg/head/year in 2001 to 20.1 kg/head/year in 2013 (35), mostly in the form of fresh pork. Vietnam has been importing breeding hogs from the U.S.A. since 1996 to enhance its swine gene-pool. On average, 553 breeding hogs were imported each year from the U.S.A. during the 2010s, with a peak in 2014–2015 (Fig S2) (36). The H1-*δ*1a lineage may have entered Vietnamese swine from imported breeding hogs in early 2016, similar to introduction into mainland China in the early 1990s (37).

In each of the SwIV lineages detected in Vietnam, we discovered evidence of localized circulation, as evidenced by periods of unsampled diversity and long phylogenetic branches, likely at the provincial level. Despite the limited mixing of swine populations in Vietnam before slaughter (33), the infrequent detection of persistent lineages in the central slaughterhouse suggests that the genetic diversity of SwIV in Vietnam may be high. This diversity is likely driven by factors such as livestock density and turnover. In comparison, swine in the U.S.A. are exposed to a greater degree of mixing during their lifespan as they are transported across long distances for feeding and fattening, which results in the replacement by advantageous SwIV lineages (27).

Our findings indicate that the SwIV gene pool in Vietnam is being enriched by importations from North America and other Asian countries, including a novel cluster of H1-*δ*1a (genotype 26) viruses that may pose a zoonotic threat (38). The H1-*δ*1a virus was only found during the last sampling visit and its persistence is uncertain. The repeated transmission of H1N1pdm09 viruses from humans to swine and reassortment with prevailing SwIVs also increased genetic diversity. Hence, to limit further introductions and diversification of the SwIV gene pool in Vietnam, it is important to actively monitor both local swine herds and imported swine.

## Materials and methods

### Surveillance and virus isolation

One hundred and fifty paired pig serum and nasal swab specimens were collected monthly through convenience sampling at the Van Phuc slaughterhouse, the main collective slaughterhouse in Hanoi, Northern Vietnam. Province of origin, pen number, and trader’s name were recorded for each sampled pig. Serum samples collected from 7 May 2013 to 28 August 2019 are reported in this study. Virus isolation and sequence data on swabs collected from 7 May 2013 to 19 May 2016 were reported previously (13) and virus isolation and sequence data on swabs collected from 30 June 2016 to 28 August 2019 are reported in the current study. Samples were cultured for virus isolation in Madin-Darby Canine Kidney (MDCK) cells at the National Institute of Veterinary Research laboratory in Hanoi, Vietnam using described methodology (13).

### Whole-genome sequencing and assembly

The influenza genome was amplified from RNA extracted from virus isolates using a multi-segment RT-PCR with universal primers (39, 40) and sequenced using Nextera DNA library preparation and MiSeq (PE300) sequencer (Illumina, San Diego, CA, USA), as previously described (13). Adapters and low-quality reads were removed using Trimmomatic v0.36 (41), and *de novo* assembly was performed using IVA v0.8.1 (42). The assembled contiguous sequences were mapped to the reference sequence using Geneious Prime 2019.2.1, and consensus sequences were generated by merging overlapping contigs. The generated Vietnam SwIV genome sequences and associated metadata are deposited at GISAID. Sequences generated in this study were analysed with other closely related influenza viruses in GenBank and GISAID databases, determined using a BLAST search, and including vaccine strains of human seasonal influenza H1N1. H1 clades were classified according to a phylogeny-based global nomenclature system (43).

### Phylogenetic analysis and genotypic diversity

Gene segments were aligned individually using MAFFT v7.490 (44) and ML phylogenetic trees were constructed using IQ-TREE v2.1.4 (45) based on the best-fit nucleotide substitution model determined by ModelFinder (46) with branch supports estimated using an approximate likelihood ratio (SH-like) test (47). The genotypes of SwIVs were determined by assigning each segment to specific based on the ML phylogenies and characterising the genotype based on the clade distribution of its internal segments (48). Time-scaled phylogenies calibrated by sample collection dates were inferred using the ML method in IQ-TREE v2.1.4, implementing a least-square dating algorithm (LSD) (49). The trees were visualized in FigTree v1.4.4.

### Serological assays

Ten pig sera were randomly selected from each monthly sampling visit to the same slaughterhouse (13). In total, 760 pig sera collected from May 2013 to August 2019 were tested using the hemagglutination inhibition (HAI) assay against three swine H1-subtype influenza virus strains and one swine H3-subtype influenza virus strain isolated in this study, as well as the pandemic H1N1 virus. These were A/California/04/2009 (pdm09, H1N1), A/Swine/Hanoi/7-305/2016 (H1-TR, H1N2), A/Swine/Hanoi/11-260/2019 (H1*δ*-like, H1N2), A/Swine/Hanoi/12-276/2019 (H1-*δ*1a, H1N2) and A/Swine/Hanoi/10-984/2018 (2004/05 human H3N2-origin). Viruses were grown in MDCK cells and were used as antigens. The HAI tests were carried out according to the WHO standard protocol for animal influenza diagnosis (50), with details as previously described (13).

## Acknowledgements

We gratefully acknowledge the staff from the originating laboratories responsible for obtaining the specimens and from the submitting laboratories where the genome data were generated and shared via GISAID (Supplementary Data 1). This study was supported by Contract No. 75N93021C00016 from the National Institute of Allergy and Infectious Diseases, National Institutes of Health, Department of Health and Human Services, United States. The funding bodies had no role in the design of the study and collection, analysis, and interpretation of data and writing of the manuscript.

## Disclaimers

The authors declare no conflicts of interest with material associated with this manuscript.

