## Supplementary material for "Long-term epidemiology and evolution of swine influenza viruses in Vietnam": Fig S1

**Supplementary Figure S1.** Maximum likelihood phylogeny of each segment of SwIV in Vietnam. Maximum likelihood tree was constructed with IQ-TREE (53). Colors indicate HA lineage. Viruses from outside Vietnam are shown in gray. Isolates with no HA sequence are shown in pink. Scale bars represent nucleotide substitutions per site.

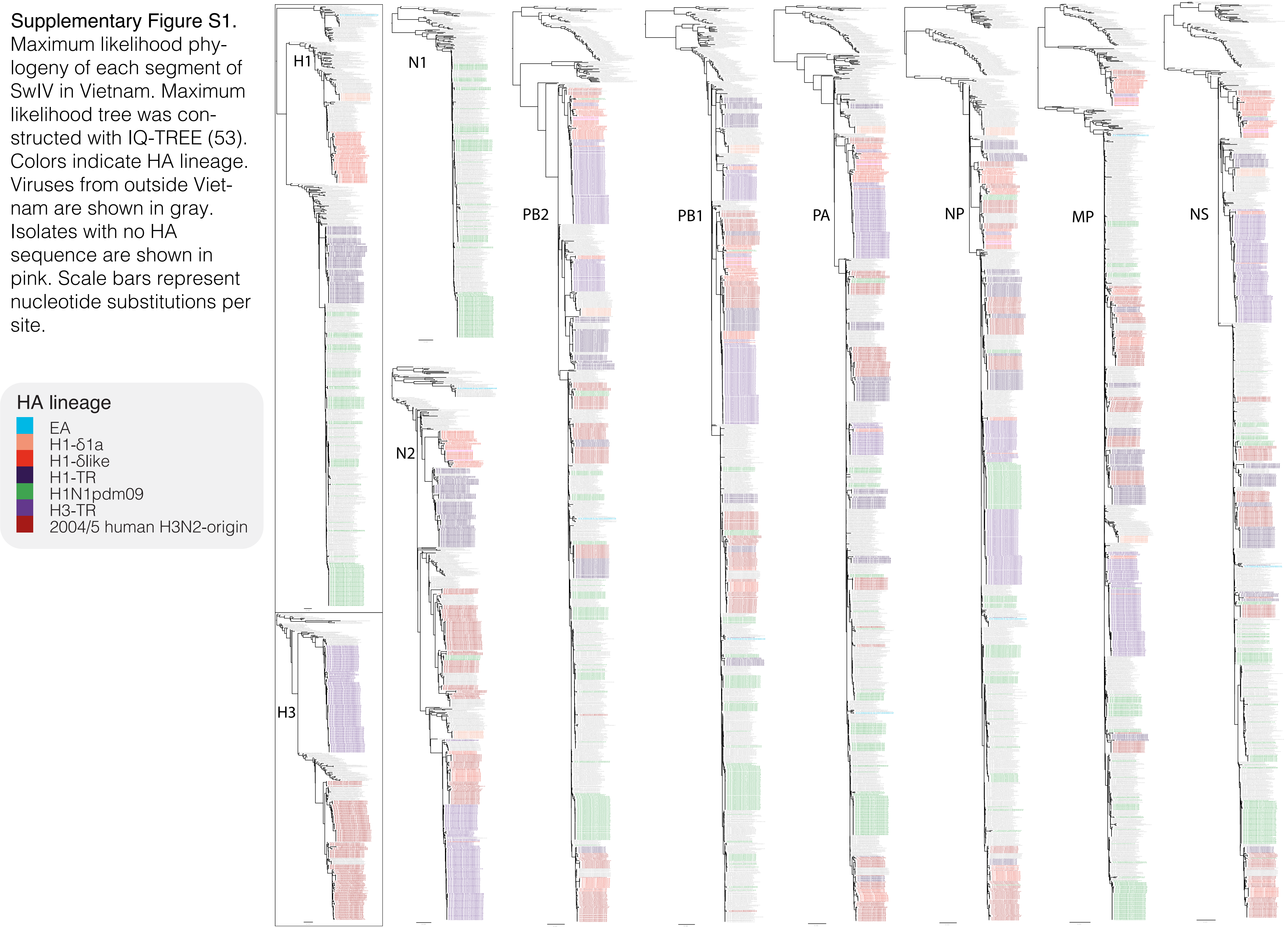
