## Supplementary figures and images for "Long-term epidemiology and evolution of swine influenza viruses in Vietnam"

### Fig S2

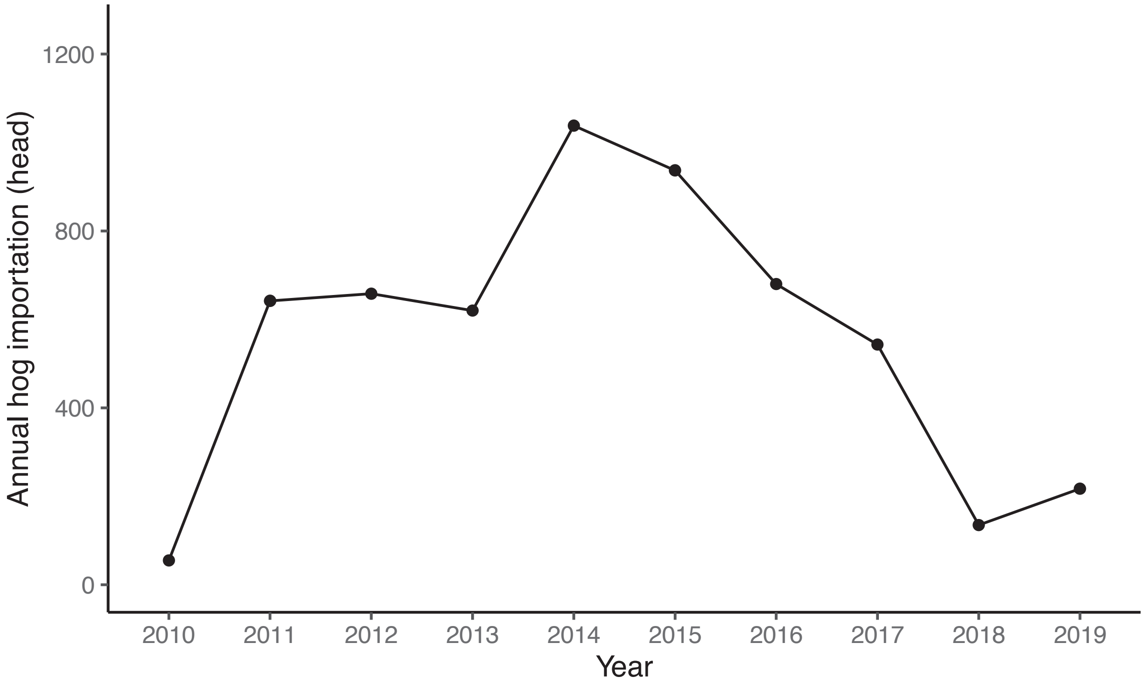


Supplementary Figure S2. Annual hog importation from U.S.A. to Vietnam in 2010–2019 (Ref. 40).
