## Supplementary material for "Long-term epidemiology and evolution of swine influenza viruses in Vietnam": Table S1

**Supplementary Table S1.** Seroprevalence of antibodies against the swine H1-$\delta$1a virus (A/swine/Hanoi/12-276/2019) in pigs in Northern Vietnam.

| **Year** | **No. of swine sera tested by HAI assay** | **No. (%) seropositive*** |
|---|---|---|
| **2013** | 80 | 0 (0) |
| **2014** | 120 | 0 (0) |
| **2015** | 120 | 0 (0) |
| **2016** | 120 | 12 (10) |
| **2017** | 120 | 8 (6.7) |
| **2018** | 120 | 4 (3.3) |
| **2019** | 80 | 12 (15) |
| **Total** | **760** | **36 (4.7)** |

* Seropositivity defined as HAI titer ≥40.
