## Supplementary material for "Long-term epidemiology and evolution of swine influenza viruses in Vietnam": Table S2

**Supplementary Table S2.** HAI titers for individual H1-$\delta$1a seropositive swine sera against other H1 SwIV lineages. Titers ≥40 are shown in bold.

| Sera ID | Sampling Date | H1-$\delta$1a A/swine/Hanoi/12-276/2019(H1N2) | H1-$\delta$-like A/swine/Hanoi/11-260/2019(H1N2) | H1N1pdm09 A/California/04/2009 (H1N1) | H1-TR A/swine/Hanoi/7-305/2016(H1N2) |
|---|---|---|---|---|---|
| **Ta_6_101M** | 24/3/2016 | **320** | 20 | <10 | <10 |
| **Ta_6_667M** | 21/7/2016 | **160** | 20 | **160** | **40** |
| **Ta_6_969M** | 21/9/2016 | **640** | **160** | **160** | **40** |
| **Ta_7_119M** | 27/10/2016 | **640** | **320** | **40** | <10 |
| **Ta_7_120M** | 27/10/2016 | **80** | <10 | 10 | <10 |
| **Ta_7_121M** | 27/10/2016 | **160** | 20 | **80** | **40** |
| **Ta_7_301M** | 25/11/2016 | **80** | <10 | **40** | <10 |
| **Ta_7_304M** | 25/11/2016 | **640** | **40** | 10 | <10 |
| **Ta_7_310M** | 25/11/2016 | **80** | 10 | <10 | <10 |
| **Ta_7_313M** | 25/11/2016 | **160** | 10 | <10 | <10 |
| **Ta_7_324M** | 25/11/2016 | **80** | **40** | **80** | **40** |
| **Ta_7_422M** | 28/12/2016 | **80** | <10 | <10 | <10 |
| **Ta_7_721M** | 23/2/2017 | **640** | **40** | **40** | <10 |
| **Ta_7_871M** | 23/3/2017 | **80** | <10 | 20 | <10 |
| **Ta_8_170M** | 25/5/2017 | **160** | <10 | 20 | <10 |
| **Ta_8_321M** | 22/6/2017 | **160** | <10 | <10 | <10 |
| **Ta_8_477M** | 26/7/2017 | **160** | <10 | **80** | 20 |
| **Ta_8_627M** | 24/8/2017 | **160** | 10 | **80** | 10 |
| **Ta_8_772M** | 28/9/2017 | **80** | <10 | **80** | 10 |
| **Ta_8_774M** | 28/9/2017 | **160** | <10 | **40** | <10 |
| **Ta_10_129M** | 22/6/2018 | **80** | **80** | <10 | <10 |
| **Ta_10_578M** | 27/9/2018 | **320** | **320** | <10 | <10 |
| **Ta_10_729M** | 30/10/2018 | **160** | **≥1280** | 20 | <10 |
| **Ta_10_875M** | 27/11/2018 | **80** | 20 | <10 | <10 |
| **Ta_11_622M** | 24/4/2019 | **160** | **40** | 10 | 10 |
| **Ta_11_631M** | 24/4/2019 | **80** | 20 | <10 | <10 |
| **Ta_11_779M** | 23/5/2019 | **80** | **160** | <10 | <10 |
| **Ta_11_928M** | 25/6/2019 | **≥1280** | **160** | **80** | 10 |
| **Ta_11_929M** | 25/6/2019 | **80** | 20 | 20 | <10 |
| **Ta_11_931M** | 25/6/2019 | **80** | 10 | 10 | <10 |
| **Ta_12_075M** | 17/7/2019 | **80** | **640** | <10 | <10 |
| **Ta_12_079M** | 17/7/2019 | **80** | 10 | <10 | <10 |

| | | | | | |
|---|---|---|---|---|---|
| **Ta_12_296M** | 28/8/2019 | **80** | 20 | 20 | <10 |
| **Ta_12_298M** | 28/8/2019 | **80** | **80** | <10 | <10 |
| **Ta_12_300M** | 28/8/2019 | **80** | <10 | 10 | <10 |
| **Ta_12_304M** | 28/8/2019 | **40** | **160** | 10 | <10 |
