## Supplementary material for "Long-term epidemiology and evolution of swine influenza viruses in Vietnam": Table S3

**Supplementary Table S3.** HAI titers for individual H1-TR seropositive swine sera against H1N1pdm09 antigen. Titers ≥40 are shown in bold.

| **Sera ID** | **Sampling Date** | **H1N1pdm09** A/California/04/2009(H1N1) | **H1-TR** A/swine/Hanoi/7-305/2016 (H1N2) |
|---|---|---|---|
| **Ta_1_121M** | 7/5/2013 | **320** | **80** |
| **Ta_1_211M** | 11/6/2013 | **160** | **40** |
| **Ta_1_637M** | 5/9/2013 | **160** | **40** |
| **Ta_1_847M** | 3/10/2013 | **320** | **80** |
| **Ta_2_276M** | 17/1/2014 | **320** | **80** |
| **Ta_2_536M** | 6/3/2014 | **80** | **40** |
| **Ta_3_234M** | 22/8/2014 | **640** | **160** |
| **Ta_4_001M** | 29/1/2015 | **320** | **80** |
| **Ta_4_417M** | 23/4/2015 | **160** | **40** |
| **Ta_5_618M** | 23/12/2015 | **80** | **40** |
| **Ta_6_667M** | 21/7/2016 | **160** | **40** |
| **Ta_6_969M** | 21/9/2016 | **160** | **40** |
| **Ta_7_121M** | 27/10/2016 | **80** | **40** |
| **Ta_7_324M** | 25/11/2016 | **80** | **40** |
| **Ta_7_426M** | 28/12/2016 | **80** | **40** |
| **Ta_8_174M** | 25/5/2017 | **80** | **40** |
| **Ta_8_622M** | 24/8/2017 | **80** | **40** |
| **Ta_9_973M** | 24/5/2018 | **80** | **40** |
